# Transcriptional changes in the mammary gland during lactation revealed by single cell sequencing of cells from human milk

**DOI:** 10.1101/2020.11.06.371443

**Authors:** Alecia-Jane Twigger, Lisa K. Engelbrecht, Karsten Bach, Isabel Schultz-Pernice, Stefania Petricca, Christina H. Scheel, Walid Khaled

**Affiliations:** Institute of Stem Cell Research, Helmholtz Zentrum München, Munich, Germany; Department of Pharmacology, University of Cambridge, Cambridge, England; Department of Dermatology, Ruhr University of Bochum, Bochum, Germany; Wellcome-MRC Cambridge Stem Cell Institute, Cambridge, CB2 0SZ

## Abstract

Findings from epidemiological studies suggest that breast cancer risk is influenced by parity in an age-dependent manner. However, human mammary tissue remodelling that takes place during pregnancy and lactation remain little understood due to the challenge of acquiring samples. Here, we present an approach to overcome this using single-cell RNA sequencing to examine viable primary mammary epithelial cells isolated from human milk compared to resting, non-lactating breast tissue. Thereby, we determined that separate to breast tissue, human milk largely contains epithelial cells belonging to the luminal lineage, as well as immune cells. Our data reveal the presence of two distinct secretory luminal cell clusters in milk which highly express luminal progenitor signatures akin to non-lactating breast tissue luminal cells. Taking advantage of the fact that both the resting and lactating mammary gland contain a luminal compartment, we focussed on comparing these transcriptomes and identified differences in mammary cell function and metabolism between these maturation states. These findings provide the basis to dissect human luminal differentiation and milk biosynthesis pathways that in the future, may be interrogated to determine how parity influences luminal cell metabolism and breast cancer risk.

## Background

The mammary gland undergoes cycles of tissue remodeling throughout a woman’s reproductive lifespan that are particularly pronounced during pregnancy, lactation and the return of the mammary gland to its resting state post involution. Determining the differentiation dynamics driving these developmental stages is not only essential to understand normal mammary gland function, but also the origins of breast cancer. The mammary gland consists of a bilayered ductal tree with an inner layer of luminal cells that mature into secretory cells during lactation, and a basal network of contractile myoepithelial cells that support transport of the milk to the nipple during milk ejection. In the human mammary gland, this ductal tree is embedded in collagen-rich, vascularized stroma containing different types of mesenchymal and immune cells. Major changes in the architecture and cellular composition of the adult mammary gland are required for the synthesis and secretion of the complex bioactive fluid that is human milk^1^. Findings in murine models suggest that these changes have a lasting impact on the mammary epithelium at an epigenetic level^2^ and lead to the generation of parity-induced cell types^3,4^. These molecular and cellular changes occurring in the mammary gland may point toward a mechanism explaining the reduced longterm breast cancer risk associated with parity^5^ and extended periods of lactation^6^.

Single-cell transcriptomic profiling of murine mammary epithelial cells has shed light on the differentiation dynamics of mammary epithelial cells. These studies described a common luminal progenitor cell in the virgin gland that gives rise to hormone responsive mature luminal cells and, in the case of pregnancy, to secretory alveolar cells^4,7,8^. Interestingly, findings from Bach et al. determined that the post-parous mammary gland contained primed parity-induced luminal progenitor cells that upregulated lactation-associated genes^4^. This is of particular interest given that luminal progenitor cells have been proposed as the cell of origin for different breast cancer subtypes^9^.

Analogous to the murine mammary gland, an emerging numbers of studies have begun to characterise human mammary subpopulations using single cell transcriptomics^10–12^ Normal mammary tissue is usually derived from aesthetic breast reductions, an invasive procedure not performed during lactation. Thus, compared to its resting state, tissue from lactating human mammary glands is difficult to obtain. Here, we show the presence of mammary epithelial cells in human milk offers an opportunity to study lactation-induced cellular changes in an entirely non-invasive manner.

We provide a systematic analysis of the single-cell transcriptome of 54,323 viable and proliferative human mammary epithelial cells either derived from milk (29,078 lactation derived mammary cells, LMC) or resting, normal breast tissue (25,245 non-lactation derived mammary cells, NMC), each taken from four age-matched donors. We found that human milk largely contains epithelial cells belonging to the luminal lineage as well as a repertoire of immune cells. Further transcriptomic analysis of the milk cells identified two distinct secretory cell types that also shared similarities with luminal progenitors, but no populations comparable to hormone responsive cells. Taken together, our data provides a comprehensive reference map and a window on the cellular dynamics that occur during human lactation which will reveal further information on the interplay between pregnancy, lactation and breast cancer.

## Results

### Cells isolated from milk or breast tissue display distinct molecular profiles

Differences in the extraction techniques required to isolate human mammary cells from either non-lactating tissue or milk are reflective of the different environments the cells are isolated from. To isolate NMCs, tissue donated from elective aesthetic mammoplasty surgery was mechanically dissected and enzymatically digested to separate epithelial fragments that could either be immediately frozen or trypsinised further to generate single cells (Fig. 1a, Supplementary Fig. 1a). On the other hand, centrifugation of freshly expressed whole milk was sufficient to isolate single LMCs from the pellet of the colloidal suspension (Fig. 1a, Supplementary Fig. 1a). Both viably isolated NMCs and LMCs could be cultured in either 2-dimensions to generate monolayer cultures or in 3-dimensional floating collagen gels to generate mammary organoids (Fig. 1b). Following isolation, single NMCs and LMCs were compared using flow cytometry and single cell-transcriptomic analysis to determine the phenotypic differences between cells derived from these different differentiation states.

**Figure 1:**
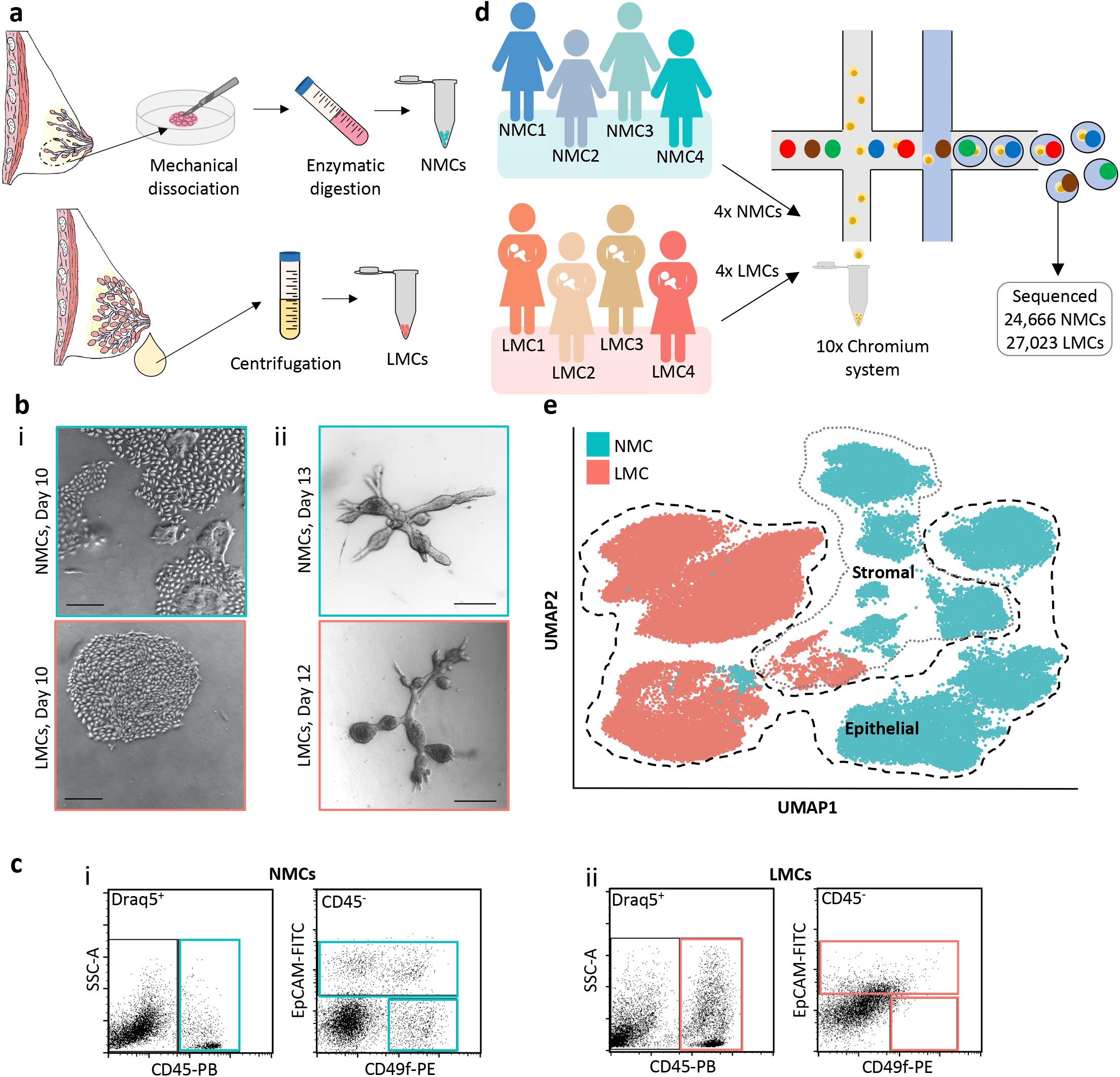
Exploring the diversity between non-lactating mammary cells (NMCs) and lactation derived mammary cells (LMCs). **a)** Cells from non-lactating tissue (above) and human milk (below) were isolated using either mechanical dissociation and enzymatic digestion or centrifugation, for downstream analysis. **b)** Mammary cells from both nonlactating breast tissue (top) or lactating milk cells (bottom) breast were cultured for either in **i)** 2D or **ii)** 3D, scale bar represents 250μm **c)** Representative flow cytometry profiles of stromal (Draq5+/CD31+ or CD45+), luminal (EpCAM+/CD49f+/-) and myoepithelial cells (EpCAM-/CD49f+) from **i)** NMCs and **ii)** LMCs. **d)** Schematic diagram for the scRNA-seq experimental set up for cell samples from 4 non-lactating participants and 4 lactating females. **e)** Uniform manifold approximation and projection (UMAP) dimensional reduction of the mammary cells reveals distinct clusters arising from NMCs and LMCs.

Using flow cytometry and previously reported markers of epithelial cell adhesion molecule (EpCAM) and integrin alpha-6 protein (CD49f)^13^, we compared the mammary subpopulation profile of gated single nucleated cells derived from either non-lactating breast tissue (n=4) or human milk (n=4). Cells isolated from non-lactating mammary tissue, contained a subset of DRAQ5^+^ single cells staining positive for CD45 (9.5-17.9%, Fig. 1c, Supplementary Fig. 1b-d). The epithelial compartment of NMCs consisted of a CD45^-^/EpCAM^-^/CD49f^+^ basal myoepithelial subpopulation (4.1-5.6%, Fig. 1c, Supplementary Fig. 1b-d) and a CD45^-^ /EpCAM^+^ luminal compartment (5.0-17.3%, Fig. 1c, Supplementary Fig. 1b-d). This luminal compartment could be further separated into CD49f^+^ luminal progenitor or CD49f^-^ mature luminal subpopulations as has been previously described^9^. Conversely, subpopulations of LMCs (n=5) were not clearly distinguishable using the same gating strategies as for the NMCs and were highly variable between participants (Fig. 1c, Supplementary Fig. 1b-d). Whilst many DRAQ5^+^ single LMCs stained positive for CD45 (4.5-43.7.9%, Fig. 1c, Supplementary Fig. 1b-c) the CD45^-^ compartment did not display a clearly distinguishable myoepithelial subpopulation (0.2-1.0%, Fig. 1c, Supplementary Fig. 1b-c) nor revealed a clear distinction between EpCAM^+^ and EpCAM^-^ cells (Fig. 1c, Supplementary Fig. 1b-c). Rather, a linear relationship between the expression of EpCAM and CD49f existed across the samples, indicating a potential loss of cell surface marker expression by these milk-derived cells. It is clear from this analysis that using only the few markers established for NMCs is insufficient to characterize subpopulations existing in LMCs.

To better define mammary cell subpopulations in human milk compared to resting breast tissue we compared 4 samples from each differentiation state using single cell RNA-sequencing. To minimize batch effects, we collected donor milk (n=4) on a single day and directly isolated the cells before loading them on a single 8-lane 10x genomics chip together with freshly dissociated cells from frozen epithelial fragments from 4 age matched tissue donors (Fig. 1d). NMCs collected were from 3 nulliparous females and one parous female (parity of 2) with ages ranging from 19-47 years old, were compared to LMCs from 3 uniparous females and one multiparous female (parity of 3) ranging from 27-43 years old (Supplementary Fig. 2a). A total of 24,666 high quality NMCs and 27,023 high quality LMCs were sequenced and retained for downstream analysis (Fig. 1d). Variance between the transcriptomic profile of single cells was examined using principle component (PC) analysis and revealed that the greatest difference (along PC1) between cells was whether they were derived from non-lactating tissue or milk (Supplementary Fig. 2b). Uniform manifold approximation and projection (UMAP), was performed to visualize the global structure of the data. In line with the PC analysis, we found little overlap between NMCs and LMCs which generally formed separate clusters (Fig. 1e, Supplementary Fig. 2c). By contrast, samples from each maturation state overlapped (Fig. 1e, Supplementary Fig. 2c). From these results, we concluded that the stark separation between LMCs and NMCs was due to their origin (milk or non-lactating tissue), rather than inter-donor variation and thus, allowed us to further probe the transcriptomic differences between the lactating and resting human mammary gland.

### Two distinct secretory clusters characterize the luminal compartment in the lactating mammary gland

Mammary cell subpopulations were identified by conducting graph clustering which revealed 5 major epithelial cell clusters across all sequenced mammary cells. Among these, 3 clusters contained cells derived exclusively from NMCs which we found to represent a single myoepithelial cluster and two luminal clusters, in agreement with previous human mammary scRNA-seq studies^10–12^. Co-expression of established myoepithelial markers encoding for transcription factor p63 (*TP63*), keratin 17 (*KRT17*), metallopeptidase CD10 (*MME*), together with contractility genes encoding for alpha smooth muscle actin (*ACTA2*), transgelin (*TAGLN*), myosin light chain kinase (*MYLK*) and tropomyosin (*TPM2*) demarked a single cluster as containing myoepithelial cells (**MY**). The two remaining NMC clusters expressed key luminal markers encoding for keratin 18 (*KRT18*) and EPCAM (*EPCAM*). Upon closer examination, one cluster resembled the hormone responsive cluster previously described^10,11^ which expressed genes encoding for hormone receptors for estrogen, progesterone and prolactin (*ESR1, PRG, PRLR* (Fig. 2a, b). The last luminal cluster resembled the previously annotated “hormone insensitive”^11^ or “secretory L1”^10^ clusters which co-expressed transcription factor *ELF5* as well as *ALDH1A3, KIT, SLPI* and *KRT23*. These markers are also characteristic of “luminal progenitor” cells in the mouse^4,7^ and hence for the purposes of this study we denote cells in this cluster as luminal progenitor (**LP**) cells (Fig. 2a, b). Thus, NMCs contain all epithelial cell subpopulations previously described in human and mouse mammary scRNA-seq studies.

**Figure 2:**
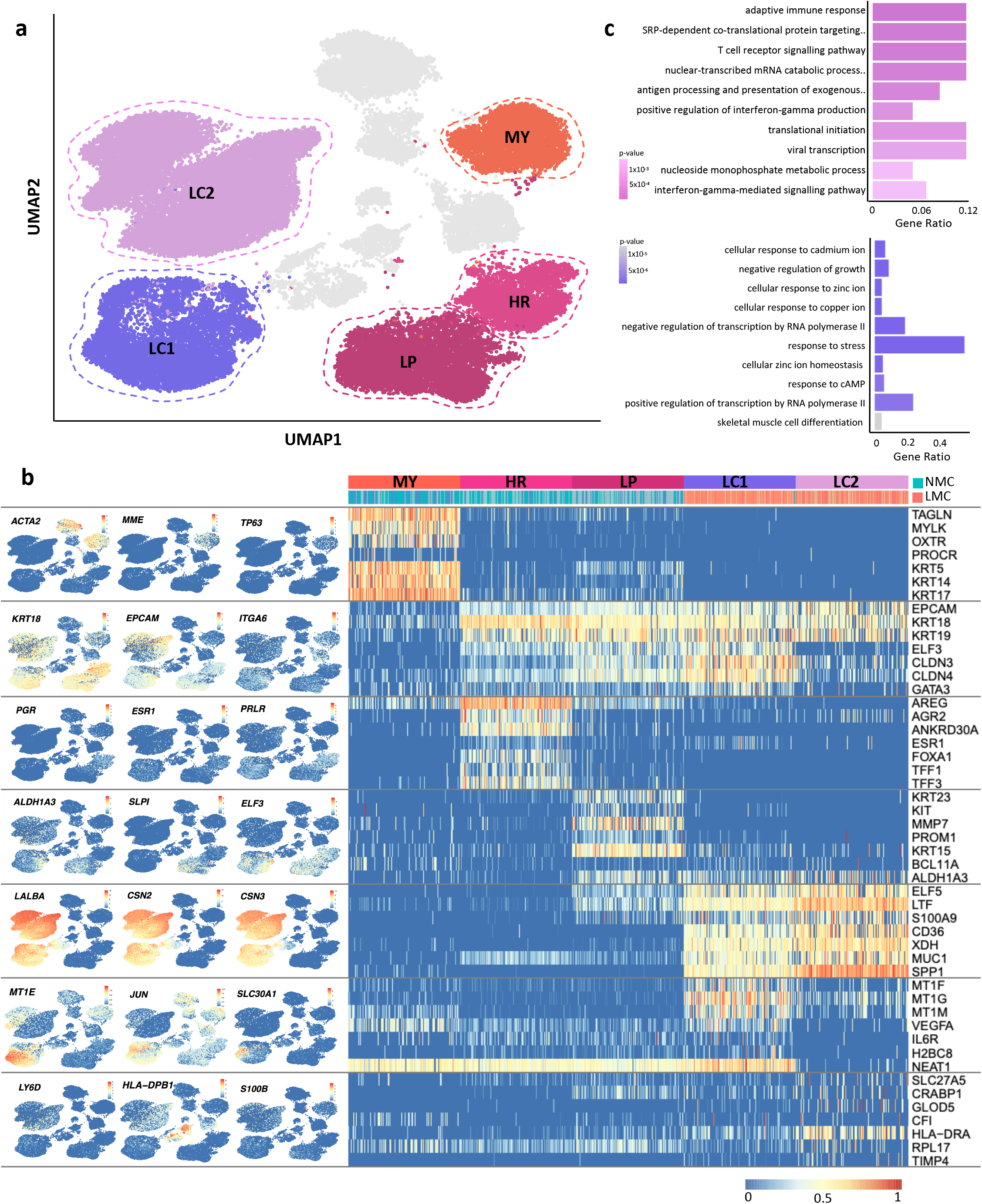
Clustering analysis of non-lactating (NMC) and lactation associated (LMCs) mammary epithelial cells reveals different subpopulations arising from different developmental stages. **a)** Five major epithelial clusters were identified in our data set consisting of a NMCs myoepithelial (MY), luminal hormone responsive (HR) and luminal progenitor (LP) cluster and LMC major luminal clusters 1 and 2 (LC1 and LC2). **b)** Examination of the various marker genes characterizing the various clusters. **c)** The top 10 biological process gene ontology pathways that were associated with genes significantly differentially expressed that were either upregulated in either LC2 (upper panel) or LC1 (lower panel) for a full list see Supplementary Table 2-3.

On the other hand, no such detailed analysis has been previously published for human milk cells, where sparse number of studies have characterized different subpopulations based on a limited number of markers. Our clustering analysis identified 2 major epithelial clusters that contained a heterogeneous contribution from all 4 LMCs donors and a very small proportion of NMCs (0.3-1.7% of total NMCs) from all (including nulliparous) donors (Fig. 1e, Supplementary Fig. 3). Due to the fact that both of these clusters co-expressed luminal markers *KRT18* and *EPCAM* as well as genes encoding the major human milk proteins *LALBA, CSN2* and *CSN3*^14^, we designated these clusters as secretory luminal clusters 1 and 2 (**LC1** and **LC2**, Fig. 2a). Cells within these clusters were found to highly express other secretory genes encoding milk fat globule membrane associated proteins xanthine hydrogenase (*XDH*), CD36 (*CD36*) and mucin-1 (*MUC1*)^15^ which are integral to milk fat secretion (Fig. 2b). Within our analysis we did not identify cells with gene expression profiles characteristic of other epithelial cell types (such as myoepithelial, stem cells) or mesenchymal stem cells (see Fig. 2b, Supplementary Fig. 4) which have been described previously^16^.

To better understand secretory luminal LMC heterogeneity, we compared **LC1** to **LC2** and found a total of 2191 genes significantly (false discovery rate corrected p-value, FDR < 0.001) differentially expressed, where 1,539 genes were found to be higher in **LC1** and 652 genes were found to be higher in **LC2** (Supplementary Fig. 3c). Significant genes upregulated in **LC1** or **LC2** were ordered according to their fold change differences and gene set enrichment analysis for gene ontology (GO) biological process terms was performed on the top 10% (153 and 65 genes, respectively). High expression of multiple major histocompatibility complex class II genes by cells within **LC2** (Fig. 2b-c, **Supplementary Table 1, 2**) suggests that these epithelial cells may conduct antigen presentation, as has been seen in other epithelial cells within the gastrointestinal and respiratory tracts^17^. In addition, an upregulation of ribosomal protein encoding genes suggests that **LC2** are more transcriptionally active than **LC1** cells (Fig. 2b-c, **Supplementary Table 1, 2**). Also among the top significant genes upregulated in **LC2** were luminal cell marker lymphocyte antigen 6 family member D (*LY6D*)^7^, fatty acid transporter solute carrier family 27 member 5 (*SLC27A5*) involved in bovine lactation^18^ and S100 calcium binding protein B (*S100B*) commonly expressed in human milk^19^ (Fig. 2b, **Supplementary Table 1**). In contrast, many DEGs upregulated in **LC1** were metallothionein genes that significantly mapped to metal ion associated GO terms as well as terms indicating that cells within this cluster may be stressed (see Fig. 2b-c and **Supplementary Table 1, 3** for a full list). Metallothionein genes which have been widely studied in the mammary gland due to their affiliation with breast cancer^20^, together with zinc transporter 1 (solute carrier family 30 member 1, *SLC30A1*) were among the genes found to be significantly differentially expressed and were associated with GO terms related to metal ion transport and homeostasis (Fig. 2b-c, **Supplementary Table 1, 3**). The term “response to stress” also had many **LC1** genes significantly associated with it such as the metallothionein genes and Jun proto-oncogene AP-1 transcription factor subunit (*JUN*), potentially indicating that cells within this cluster may have metabolic profiles that are more susceptible to cellular stress (Fig. 2b-c, **Supplementary Table 1, 3**). Other genes found to be among the top significantly upregulated in **LC1** were vascular endothelial growth factor A (*VEGFA*) found to promote vascularization of the mammary gland during lactation^21^ and long non-coding RNA *NEAT1*, found to be essential for lactation^22^ (Fig. 2b-c, **Supplementary Table 1, 3**). Overall, our data revealed that milk contains two distinct secretory cell populations that both highly express lactation-associated genes as well as gene expression profiles characteristic for each cell type that appear to display disparate immunomodulatory and metal ion transportation functions.

### Investigating non-lactating and human milk cell stromal heterogeneity

The remaining four major clusters identified in this study mapped to stromal subtypes predominantly from NMC’s with one major cluster also including LMCs. In agreement with previous mammary scRNA-seq studies^11^, we identified *GJPA4*^+^ (encoding gap junction protein alpha 4) vascular accessory (**VA**) *PECAM1*^+^ (encoding CD31) endothelial cells (**EN**) and *DCN^+^/LUM^+^/COL1A1*^+^ fibroblasts (**FB**) within all NMC samples (Fig. 3a, b). Unsurprisingly, no vascular accessory, endothelial or fibroblast lineage cells were isolated from any milk samples, however all samples from both LMC and NMCs contained cells belonging to the *PTPRC^+^* (encoding CD45) immune (**IM**) cluster. To better determine the different subtypes of immune cells isolated from LMCs and NMCs, we performed subclustering analysis on the cells and annotated them according to the expression of canonical immune subpopulation markers^23^. Thus, we identified 7 subclusters consisting of either myeloid or lymphocytic lineage hematopoietic cells from NMC or LMCs (Fig. 3b-c, Supplementary Fig. 2). Two NMC subclusters contained *CD68^+^/FCER1G^+^/CD14^+^* myeloid lineage cells consisting of *CD163^+^/MSR1^+^* macrophages, (where one cluster highly expressed macrophage marker *C1QB*) and *FCGR3A^+^/ITGAX^+^/CD33^+^* monocytes and neutrophils (Fig. 3c, Supplementary Fig. 5).

**Figure 3:**
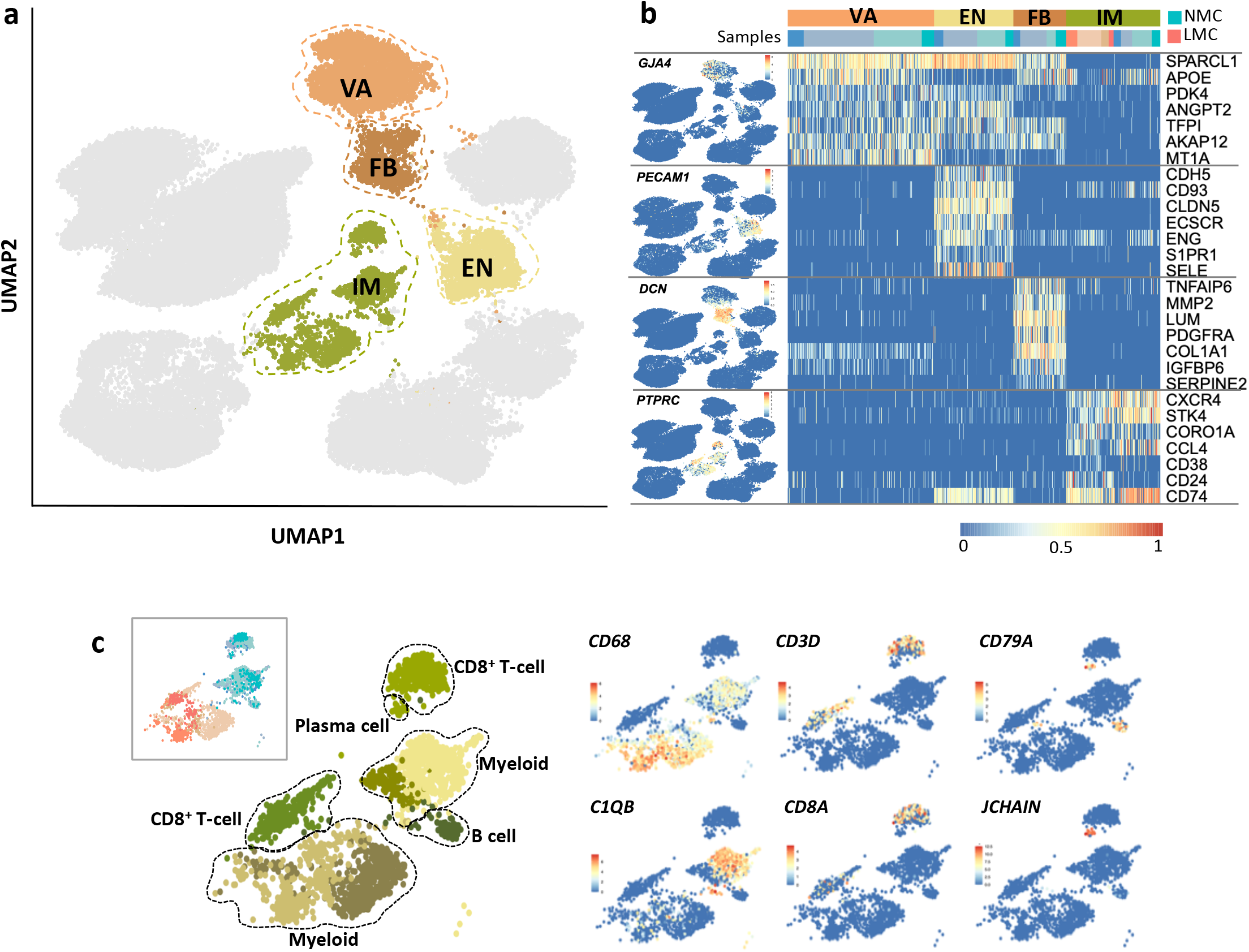
Investigation of the stromal compartment of non-lactating and lactating mammary epithelial cells. **a)** Stromal cells were classified into vascular accessory (VA), endothelial (EN), fibroblasts (FB) and immune (IM) cells. **b)** Canonical stromal markers were used to classify the different stromal subtypes where LMC only contained IM cells. **c)** Subsetting and re-clustering of IM cells revealed that both myeloid and lymphocytic lineages were sequenced from both NMC and LMCs.

A separate cluster of lymphocytic cells was identified for NMCs containing CD4+ T-cells (*CD4*^+^, *IL7R*^+^), CD8+ T-cells (*CD8A*^+^) and cytotoxic T-cells or natural killer cells expressing genes encoding granzymes A, B, H and granulysin (*GZMA, GZMB, GZMH* and *GNLY*) (Supplementary Fig. 5). NLCs also contained a further subcluster of B-cells (*CD79A^+^/MS4A1*^+^, Fig. 3c, Supplementary Fig. 5) with a small subset of *JCHAIN*^+^ plasma B cells (Fig. 3c, Supplementary Fig. 5). In comparison, LMCs contained two clusters resembling the NMC myeloid clusters and another cluster similar to the NMC lymphocytic cluster, but no B-cell cluster. We noted that LMC derived immune cells from all 3 clusters also expressed milk protein genes such as *LALBA, CSN2* and *CSN3* (Fig. 2b, 3a). Uptake of extracellular vesicles (EVs) by both myeloid and lymphocytic lineage immune cells has been previously noted to facilitate cellular communication between immune cells^24^. Finding milk transcript expression in both lymphocytic and myeloid lineage milk cells is most likely suggestive of the cells endocytosing surrounding EVs/milk fat globules containing secretory epithelial cell mRNA as part of a signaling process rather than lymphocytes acquiring macrophage phagocytosis properties. It is also possible that some of the milk protein transcripts are derived from ambient mRNA although the overall expression is high. Together this analysis finds that unlike for the epithelial cell clusters, LMCs mirror the NMC immune cell populations (except for B cells), where differences in milk protein expression might indicate mechanisms of cellular communication between mammary epithelium and immune cells during lactation. These data also highlight that similar cell types from these different sources (i.e. tissue vs. milk) display similar transcriptomes, despite the stark differences in cell-isolation protocols, suggesting that the transcript differences we observe in the epithelium compartment indeed reflect the differentiation stage.

### Exploring differences between luminal HMCs and non-lactating luminal progenitors

One major question arising from our data relates to the cell of origin of the LMC luminal cells. Which luminal population in the non-lactating, resting gland do they resemble most and are likely arise from? To address this, we examined non-lactating cell signatures^25^ derived from sorted mammary cell subpopulations^9^ across all clusters including LMCs. The purpose of this analysis was to take an unbiased approach to examine previously derived and curated gene signatures^25^ that were exclusively upregulated in hormone responsive luminal cells (n=163), luminal progenitors (n=162 genes), basal (n=125) or stromal cells (n=384) (**Supplementary Table 4**). Using these signatures, we calculated the combined expression level of these genes resulting in a subpopulation score that was visualized using violin plots for each cluster (Fig. 4a). Reassuringly, each subpopulation signature was found to display the highest expression within our data set in the cluster(s) that we had independently assigned to the same subpopulation identity. Accordingly, the hormone responsive mature luminal signature was found to be highest in **HR** cells (Fig. 4ai), the luminal progenitor score was highest in **LP** cells (Fig. 4aii), the myoepithelial score was highest in **MY** cells (Fig. 4aiii) and the stromal score was highest across a number of stromal clusters (Fig. 4aiv). These findings highlight the robustness of these gene signatures which include hundreds of markers and can be translated across bulk or single cell RNA-sequencing studies. This analysis revealed that both **LC1** and more so **LC2**, displayed the highest enrichment of the luminal progenitor score, thus displaying a similar expression pattern to NMCs within the **LP** cluster (Fig. 4a). These findings imply that secretory luminal cells from milk are most similar to luminal progenitor cells, suggesting that **LC1** and **LC2** are derived from **LP** cells. These results are in agreement with findings in the mouse^4,7,8^.

**Figure 4:**
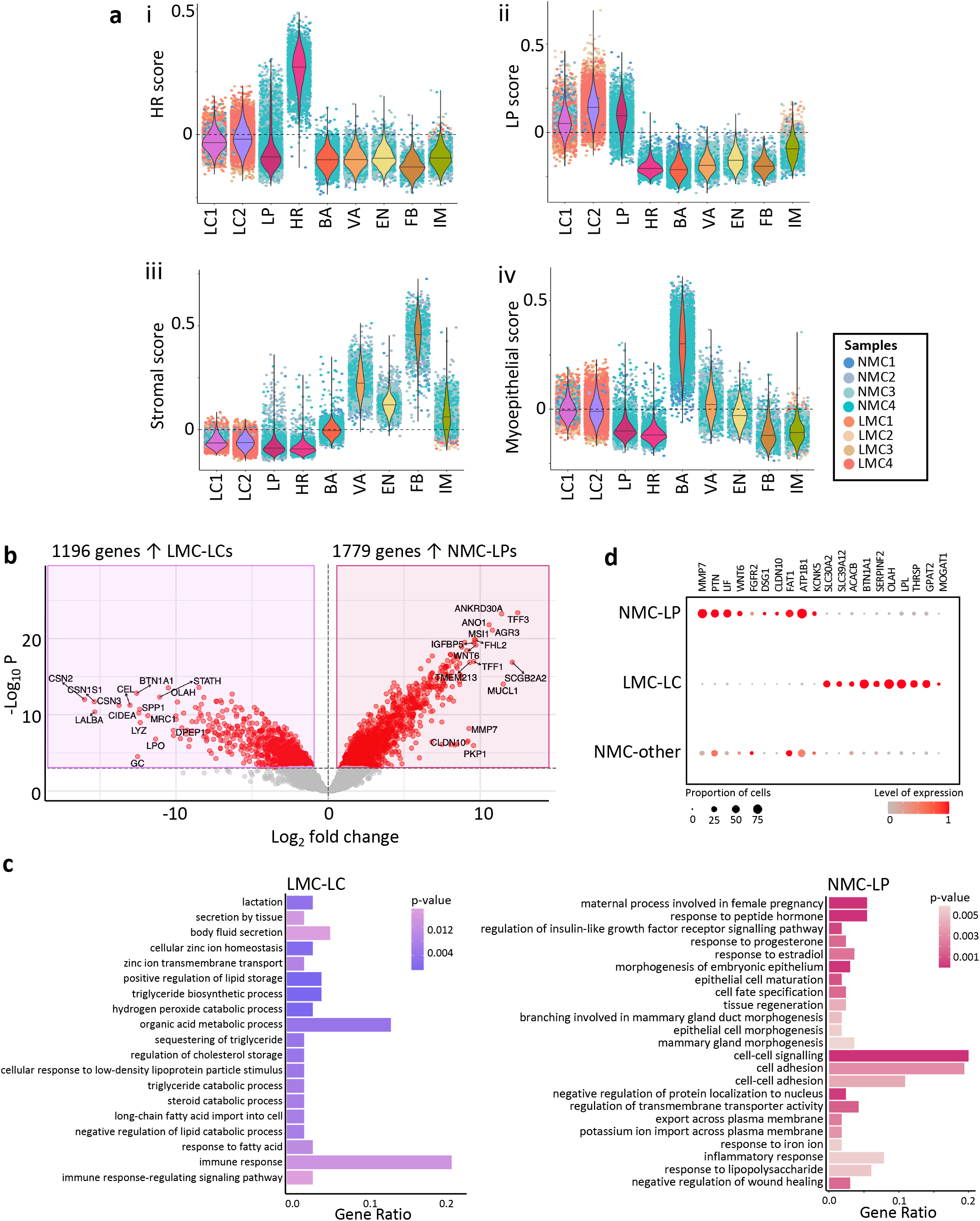
Comparing lactation derived mammary cell (LMC) luminal clusters (LCs) with all other non-lactating mammary cell (NMC) types reveals similarity to nonlactating luminal progenitor (LP) cells. **a)** Violin plots of the mammary cell scores for either **i)** hormone responsive (HR, mature luminal), **ii)** luminal progenitor (LP) **iii)** stromal or **iv)** myoepithelial cells across the major cell clusters identified in this study. **b)** Differential gene expression analysis revealed 1196 genes highly expressed in LC LMCs compared to 1779 genes more highly expressed in LP NMCs as displayed by a volcano plot (for a full list see Table S2). **c)** Important biological process gene ontology pathways that were annotated to by either genes significantly differentially expressed and upregulated in either LC (left) or LP (right) for a full list see Table S3-4. **d)** Key LP (left) or LC (right) genes expressed in NMC-LP, LMC-LC, or all other NMC clusters, colours represent overall normalised gene expression and size equals cell proportions.

Next we compared **LP** cells from NMCs with both LMC secretory luminal clusters **LC1** and **LC2** (collectively referred to as **LC**) using differential gene expression analysis. Thus, 1196 genes that were found to be significantly upregulated in **LC** clusters and 1779 genes that were significantly upregulated in **LP** cells (p <0.001, Fig. 4b, **Supplementary Table 5**). After ranking the significant differentially expressed genes by their fold change, we took a closer look at the top 10% (178 genes for LP and 119 genes for LC) and the GO biological process terms they were associated with.

Overall, GO terms associated with upregulated genes in **LC** LMCs were collectively related to fatty acid metabolism/storage, zinc transport, secretion and immune response (see Fig. 4ci and **Supplementary Table 5, 6** for detailed GO terms). Genes associated with similar terms have been previously found to be upregulated during lactation in mouse and human milk studies when comparing between different stages of development^4,26,27^ but have not been previously examined in lactating human mammary epithelial cells. Identification of genes such as these that are likely to be involved in milk biosynthesis during human lactation provide insight into the normal metabolism of the mammary gland and which in future may be used as a comparison to aberrant mammary gland metabolism of breast cancer cells.

On the other hand, genes found to be expressed at higher levels in **LP** NMCs were related to GO terms that could be broadly associated with changes to epithelial cell state, hormone response, cell trafficking, inflammation or cell adhesion (see Fig. 4cii and **Supplementary Table 7** for a full list of related GO terms). Interestingly, we find that many genes encoding chemokines such as interleukin 6 (*IL6*) and C-X-C motif chemokine ligands 1 and 6 (*CXCL1* and *CXCL6*) were downregulated in **LC** LMCs and contained in GO terms such as “inflammatory response” (Fig. 4b, cii, **Supplementary Table 5, 7**), which may reflect differences in signaling to the microenvironment in the resting compared to lactating condition. Indeed, differences between how cells interact with one another between the resting and lactating condition was highlighted by many GO terms, such as “cell adhesion” which contained genes such as desmoglein 1 (*DSG1*), tight junction protein claudin 10 (*CLDN10*)^28^ expressed in mammary cells and FAT atypical cadherin 1 (*FAT1*) which were significantly downregulated in **LC** LMCs (Fig. 4b, cii, **Supplementary Table 5, 7**). These data suggest that LMCs in milk may have downregulated many of their cell-cell adhesion molecules, either through an active process during lactation or due to being in suspension. Taken together, these data demonstrate the utility of mammary cells from milk to provide insights into the functional mammary gland and determine genes integral to normal maturation and milk secretion which cannot be identified through animal studies.

## Discussion

To better understand how the mammary gland changes during lactation, we performed flow cytometric and single cell RNA-sequencing to compare for the first-time differences in the mammary cell composition and signaling of cells isolated from non-lactating tissue and human milk. As expected, we found that the non-lactating tissue contained the full complement of mammary epithelial cells including luminal progenitor, luminal hormone responsive and myoepithelial basal cells as well as stromal cells consisting of vascular accessory, endothelial, fibroblasts and hematopoietic cells^10–12,23^. Conversely, our analysis of LMCs isolated from human milk did not reveal the same cellular subpopulations found in NMCs, nor contain the full variety of milk cell subpopulations that have been previously determined using analysis with limited numbers of markers^14,16,29–34^. Instead, using thousands of markers per cell, we identified through scRNA-seq, two unique secretory luminal clusters derived from all LMC samples that were distinct from NLC epithelial clusters, as well as hematopoietic LMCs from both myeloid and lymphocytic lineages. Strikingly, however, we did not find any milk derived luminal cell subpopulation that resembled hormone responsive NMCs, which is in line with previous mouse studies^4,35^. This raises the question of how cells enter into human milk. Cells are likely to be secreted into milk by firstly detaching (resulting in a downregulation of adhesion markers), entering either the lumen of the alveoli or ducts and being transported together with other secretions through the nipple to be delivered to the breastfeeding infant. Considering the cellular organization of the mammary gland^1^, it is therefore conceivable that cell subpopulations that exist distal to these sites (such as fibroblasts or myoepithelial cells) do not enter the milk, compared to the more proximally localized luminal cells.

Curiously, unlike mammary studies conducted in mice^4,7^, we did not identify a single luminal secretory population but rather two subpopulations in our human samples (**LC1** and **LC2**). Whilst both clearly display known, as well as novel secretory genes, we find that additionally **LC2** cells expressed high levels of immunomodulatory and antigen presenting genes not previously associated with mammary luminal cells. The secreted proteins, through delivery to the infant may play a role in protection of the vulnerable infant or indeed provide a mechanism for training the adaptive immune system of the infant^36^. This finding raises the question of why cells enter into human milk. Some studies suggest that LMCs may enter into the milk for delivery to the infant and subsequently infiltrate into different organs for the benefit of the infant^36–38^, whilst others postulate that they enter due to a natural shedding process^39^. Future studies should examine how cells enter into the milk to better understand their potential function. Within this study, we utilized cells in milk to compare between NMCs and LMCs to gain insight into the potential epithelial differentiation trajectories that occur over mammary maturation.

Comparisons between the **LC** LMCs and **LP** NMCs did identify differentially expressed genes which provide clues into the maturation pathways modified during lactation as well as potential genes involved in milk component biosynthesis pathways that may be in future targeted in the case of low milk production. Furthermore, metabolic profiles of the luminal cells were found to be very different between NMC and LMCs, where greater participant heterogeneity in secretory luminal cells suggests that inter-individual differences in cellular metabolism may be amplified during lactation. Using functional, milk derived luminal cells instead of non-lactating breast tissue cells to compare to breast cancer cells may be instrumental to understanding tumor cell metabolism and aid in identifying therapeutic targets against pro-survival signaling^40^. Further, examining luminal progenitor-like cells from human milk may be useful as a pre-clinical tool for early detection of disease. In the future, more indepth characterization of milk cells cultured as organoids *in vitro* will shed light on their full utility. Together our study demonstrates the power of comparing mammary cells isolated from different stages of human mammary gland maturation and illustrates the luminal lineage remodeling that occurs during lactation.

## Methods

### Tissue collection and non-lactating cell isolation

Non-lactating human breast tissue was donated by participants undergoing elective aesthetic reduction mammoplasty at the Nymphenburg Clinic for Plastic and Aesthetic Surgery in accordance with the regulations of the ethics committee of the Ludwig-Maximilian University, Munich, Germany (proposal 397-12). Limited demographic information on the participants was provided which included age and parity of the participant (Supplementary Fig. 2A). Single cell suspensions of mammary cells were generated using an adaptation of a previously described protocol^41^ however in this case using a fast tissue digestion protocol. Briefly, the freshly collected mammary tissue was collected and minced using scalpels into smaller than 2-3mm^3^ pieces. 20mL of minced tissue in digestion buffer (DMEM/F12 w/o phenol red, 2% w/v BSA, 10 mM HEPES, 2 mM glutamine, 100 U/ml penicillinstreptomycin) supplemented with 1 μg/mL insulin (Sigma, I6634) was added together with 800 U/mL collagenase (Sigma, C9407) and 100 U/mL hyaluronidase (Sigma, H3506) and made up to the 25mL mark. Falcon tubes were then sealed with parafilm to ensure they were airtight and mixed at 100rpm at 37°C for 3-4 hours. Following digestion, the resulting dissociated organoid fragments were washed twice in washing buffer (DMEM/F12 w/o phenol red, 10mM HEPES, 2mM glutamine, 100 U/ml penicillin-streptomycin) and pelleted by 300g centrifugation for 5 minutes. Once the organoids were isolated, they were cryopreserved in 50% washing buffer, 40% fetal calf serum and 10% DMSO and stored in liquid nitrogen until required. When required, organoid fragments were gently defrosted in a 37°C water bath for approximately 5 minutes before being treated with Trypsin and dispase (Life Technology) to yield a highly viable single cell suspension.

### Human milk cell isolation

Human milk donors were recruited from Pippagina English Prenatal and Postnatal classes or through the Helmholtz Zentrum München in accordance with regulations of the ethics committee of the Ludwig-Maximilian University, Munich, Germany (proposal 17–715). Participants provided written informed consent and filled out a detailed questionnaire to provide demographic information. Briefly, human milk was freshly collected using either a provided double electric breast pump (Medela, Symphony) or participants personal pump, under aseptic conditions. Milk collections were obtained either within the lactation room at the Helmholtz Centre Munich, at the participants homes or post-partum educational classes depending on the preferences of the participant. Fresh milk samples were immediately transported on ice to the laboratory and processed as soon as possible (< 2 hours after collection). Briefly, human milk cells were isolated by diluting milk samples in an equal volume of sterile phosphate-buffered saline (PBS) (ThermoFisher Scientific, Waltham, U.S.) and centrifuged at 870 g for 20 minutes at 20°C in a Rotanta 460R centrifuge (Hettich, Tuttlingen, Germany). The pellet was washed by removing the supernatant and resuspending in 5-10 mL of cold PBS before transferring the sample to a new 15 mL tube (Corning, Corning, U.S.) and centrifuging at 490 g for 5 minutes at 4°C. Following a second washing step, 100-550 μL of mammary epithelial cell growth medium (MECGM) (PromoCell, Heidelberg, Germany) was added to the human milk cell aggregations according to the pellet size. To examine the cells more closely, DRAQ5™ (62254, ThermoFisher Scientific, Waltham, U.S.) and Nile red (N3013-100MG, Merck, Darmstadt, Germany) were added to a final concentration of 0.4 μg/mL (1 mM) and 0.1 μg/mL respectively and the cells incubated for a further 5 minutes in the dark. Cells were then loaded onto a Neubauer Improved counting chamber and examined on an immunofluorescence microscope. Subsequently, single cells were either frozen or used immediately (in the case of scRNA-sequencing).

### Cell culture

Both NMC and LMCs were cultured in 2D and 3D using previously described methods^42^. Briefly, single cells were mixed with MECGM supplemented with 1% pen/strep (Invitrogen), 0.5% FCS (Pan Biotech), 3 μM Y-27632 (Biomol) and 10 μM forskolin (Biomol) and seeded onto polystyrene cell culture plates for 2D culture. After an establishment period of 5 days medium was changed to MECGM supplemented with 1% pen/strep and 10 μM forskolin. For 3D culture, single cells were mixed with neutralizing solution and acidified rat tail collagen I (Corning) to generate collagen gels in siloxane coated 24-well plates. After allowing the gels to polymerise for an hour, supplemented media (as above) was added on top of the gels which were then gently encircled to generate floating collagen gels. Similar to 2D culture, after an establishment period of 5 days the media on the floating collagen gels was changed to MECGM supplemented with pen/strep and forskolin only.

### Flow cytometry

Flow cytometry was employed to determine the similarity of expression of mammary markers between LMC and NMCs. Cells were stained with DRAQ5™ (ThermoFisher Scientific, Waltham, U.S.) to a final concentration 1μM, CD45-PB (dilution of 1:100), EpCAM-FITC (dilution of 1:10) and CD49f-PE (dilution of 1:20). After incubation, stained MESs were diluted in MECGM and filtered through 35 μm cell strainer caps of round-bottom tubes (Corning, Corning, U.S.). Small volumes of cells from each sample were mixed and used as comparison and control. Flow cytometry was performed using a FACSAria™ III cell sorter (BD Biosciences, Franklin Lakes, U.S.) with a 100 nm nozzle in combination with FACSDiva™ 6.0 Software. Laser settings were adjusted using unstained and single stain controls. Obtained data was analysed using the FlowJo_V10 Software (FlowJo LLC, Ashland, U.S.).

### Library preparation, sequencing and data processing

Library preparation was performed 10X Chromium single-cell kit using version 3 chemistry according to the instructions in the kit. The libraries were then pooled and sequenced on a NovaSeq6000 S2. Read processing was performed using the 10X Genomics workflow using the Cell Ranger Single-Cell Suite version 3.0.2. Samples were demultiplexed using barcode assignment and unique molecular identifier (UMI) quantification. The reads were aligned to the hg19 reference genome using the pre-built annotation package obtained from the 10X Genomics website (https://support.10xgenomics.com/single-cell-gene-expression/software/pipelines/latest/advanced/references). All lanes per sample were processed using the ‘cell ranger count’ function. The output from different lanes were then aggregated using ‘cellranger aggr’ with -normalise set to ‘none’.

### Quality control and data pre-processing

All downstream analysis was conducted using R version 4.0.0. Barcodes identified as containing low counts of UMIs likely resulting from ambient RNA were removed using the function “emptyDroplets” from the *DropletUtils* package^43^. Barcodes arising from single droplets were then filtered to ensure that cleaned barcodes contained at least 1000 UMIs and that the percentage of mitochondrial genes compared to overall annotated genes were not higher than 2 x the median absolute deviation (MAD). Overall, 13,102 cells were obtained for LMC1, 2,172 cells were obtained from LMC2, 5,900 from LMC3 and 5,849 cells from LMC4. In addition, 5,339 cells were profiled from NMC1, 6,943 cells from NMC2, 6,699 from NMC3 and 5,685 cells from NMC4. Following filtering, cleaned barcodes were normalized and log-transformed using the “computeSumFactors” from *scran* version 1.16.0 and “logNormCounts” from the *scater*^44^ package version 1.16.1. Principal component analysis was then performed on the normalized log-transformed counts using *PCAtools* version 2.0.0 followed by uniform manifold approximation and projection graphing (*umap* version 0.2.6.0) and Louvain clustering using *scran*^45^. Overall 15 clusters were identified, where six subclusters were found within the two major luminal clusters **LC1** and **LC2** which upon closer examination revealed that the donor contribution to each of the original six clusters was disproportionate (Supplementary Fig. 3a), where clustering methods may have segregated cells based on biological discrepancies between cell transcriptomic profile rather than LMC subtype heterogeneity. We noted through dendrogram analysis that these clusters were ordered into two major milk cell clusters that were separated in the same way as the UMAP and hence considered only these two major clusters for downstream analysis and interpretation (Supplementary Fig. 3a). In addition, three clusters of immune cells containing both LMC and NMCs were originally identified but represented as one cluster in initial analysis. After performing subclustering analysis on all immune cells we identified seven distinct subclusters which could be assigned to different cell subtypes based on gene expression profiles. Plots were generated using either *ggplot2* or *pheatmap* packages with custom colours generated by the *RcolourBrewer* package.

### Differential gene expression analysis

Differentially expressed genes (DEGs) were identified between subsetted groups by firstly generating pseudobulk samples using “aggregateAcrossCells” function in the *scater* package. *edgeR* version 3.30.3 was used to compute differentially expressed genes between groups by first calculating outlier genes using the “isOutlier” function, filtering by expression using “filterByExpr”, scaling library size using “calcNormFactors”. Next a model matrix was generated “model.matrix” and the common, trended and tagwise negative binomial dispersions of the genes was calculated using “estimateDisp”. Quasi-likelihood negative binomial generalized log-linear models were fitted using “glmQLFit” and “glmQLFTest”. False discovery rate corrections were applied to the resulting p-values using the Benjamini-Hochberg method. To visualize the differentially expressed genes volcano plots were generated using the *EnhancedVolcano* package the false discovery rate corrected p-value cut off here is p < 0.001 (1×10^-4^). Significant genes (p < 0.001) were ranked according to their fold change and the top 10% of the positive (upregulated) or bottom 10% of the negative (downregulated) genes had gene set enrichment analysis performed on them using the “weight01” algorithm and “fisher” statistic using “runTest” in the “getSigGroups” function from *topGO* package. “GenTable” was used to generate a table with the top 50 biological process GO terms. Plots of selected GO terms were generated using *ggplot2*, plotting the resulting p-value from the classic fisher test and gene ratio which is the number of significant genes for the term divided the total number of significant genes used in the gene enrichment test.

### Mammary cell signature score comparisons

Mammary signatures^25^ from previously published data^9^ from luminal progenitor, mature luminal, myoepithelial and stomal cells were investigated in our data using the “AddModuleScore” function from the *Seurat* package^46^ version 3.1.5. For each test the overall expression of the genes/features from each signature was calculated after subtracting 20 randomly selected genes (from the same bin as the signature features) as a control feature per cell. The resulting signature score is unitless but is indicative of signature enrichment per cell which was then compared between clusters. Few genes in the published score were^45^ not found in our data set and these have been reported in **Supplementary Table 4**.

### Data availability

The authors declare that all data supporting the findings of this study are available within the article and its supplementary information files or from the corresponding author upon reasonable request. Submission of raw gene expression and barcode count matrices to ArrayExpress is in process. For inquiries contact authors. We will also release a user-friendly website to enable Data exploration. All computational analyses were performed in R (Version 4.0.0) using standard functions unless otherwise indicated. All Codes used will be available online at https://github.com/.

## Acknowledgements

Many thanks are extended to Dr Anika Böttcher, Dr. Michael Sterr and Ms. Ines Kunze from the Helmholtz Diabetes Center for assistance with preparing the samples for single cell RNA-sequencing. Thank you to the Helmholtz sequencing core and particularly to Ms. Sandy Loesecke for conducting the sequencing. We thank Dr. Thomas Walzthoeni for bioinformatics support provided at the Bioinformatics Core Facility, Institute of Computational Biology, Helmholtz Zentrum München. Thank you to Hilary Ganz for her assistance with isolating cells for this study. A big thank you is extended to Ms. Lynn Darbyshire from Pippagina for her continued enthusiasm and support in collecting samples for this study and especially to all the women who provided their samples, without which this research would not be possible. A.J.T is funded by the ISRHML trainee bridge fund postdoctoral fellowship, the Helmholtz Postdoctoral Fellowship and a BBSRC project grant (BB/S006745/1) to W.T.K. K.B. is funded by a Cambridge Cancer Centre PhD studentship and a CRUK Career Establishment Award (C47525/A17348) to W.T.K.

## Author Contributions

A.J.T. and C.H.S. conceptualized the study. A.J.T., L.K.E., I.S.P and S.P. collected samples and prepared samples for analysis. A.J.T. and I.S.P. performed flow cytometry and culturing experiments with assistance from L.K.E. A.J.T. performed bioinformatic analysis with assistance from K.B.. A.J.T., L.K.E., I.S.P., K.B., C.H.S., W.T.K., interpreted the data. A.J.T., C.H.S. and W.T.K. wrote the manuscript with critical input from all authors.

## Supplementary Information

**Supplementary Figure 1:**
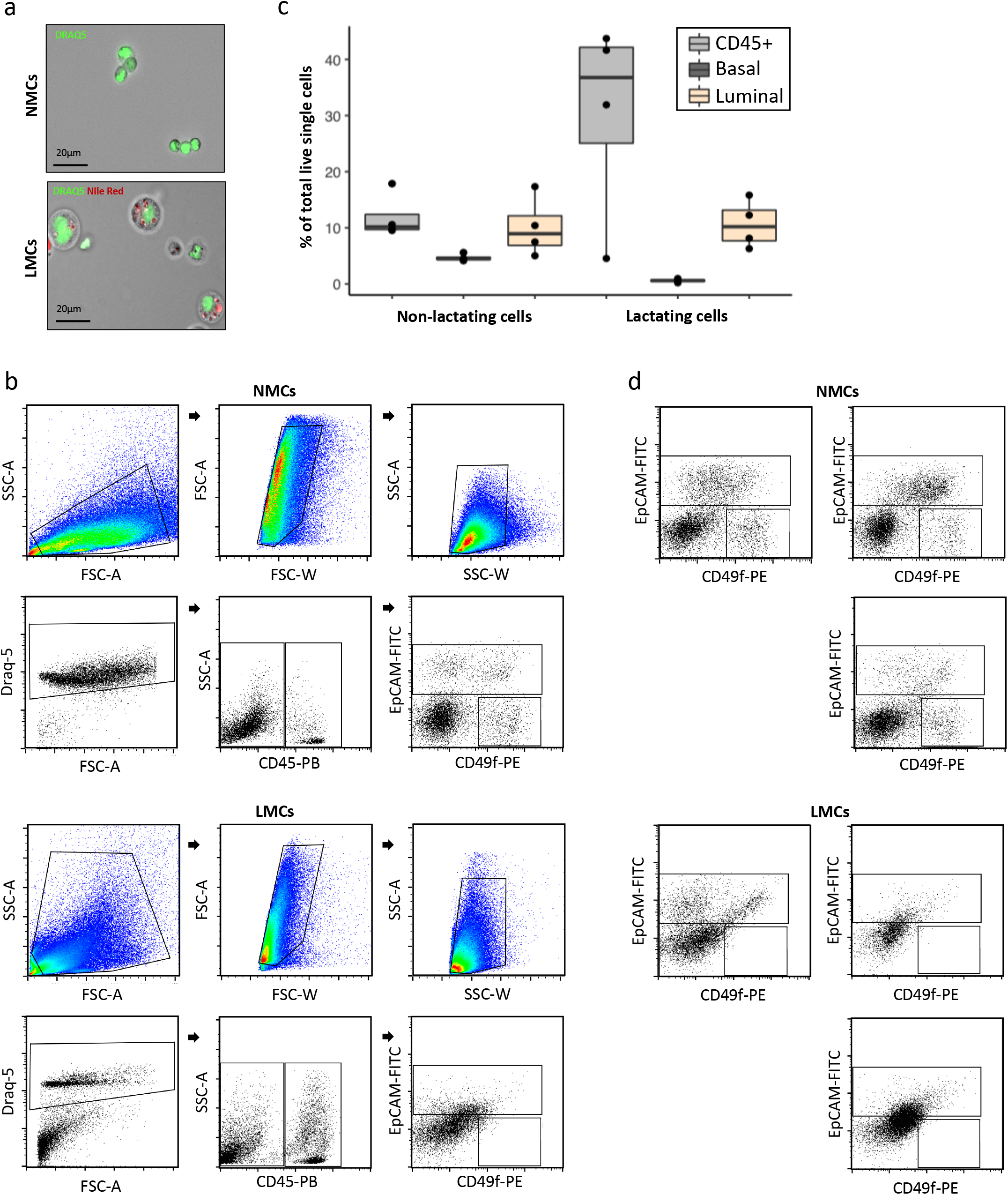
Flow cytometry plots showing full gating for non-lactating mammary cells (NMC) or lactating mammary cells (LMC) to reveal mammary subpopulations. **a)** Differences in cell morphology between NMCs (above) and LMCs (below) could be visualized using light and fluorescence microscopy using nuclear stain Draq5 and neutral lipid stain Nile red. **b)** representative full gating strategy shown for NMC (above) and LMC (below). **c)** summary of NMC (n=4) and LMC (n=4) that fall into the gates for single gated Draq5^+^ cells CD45^+^ immune cells, CD45^-^/EpCAM^-^/CD49f^+^ myoepithelial cells or CD45^-^/EpCAM^+^ luminal cells **d)** Individual plots for remaining donors showing gated epithelial populations (using EpCAM and CD49f) from NMC (above) or LMC (below).

**Supplementary Figure 2:**
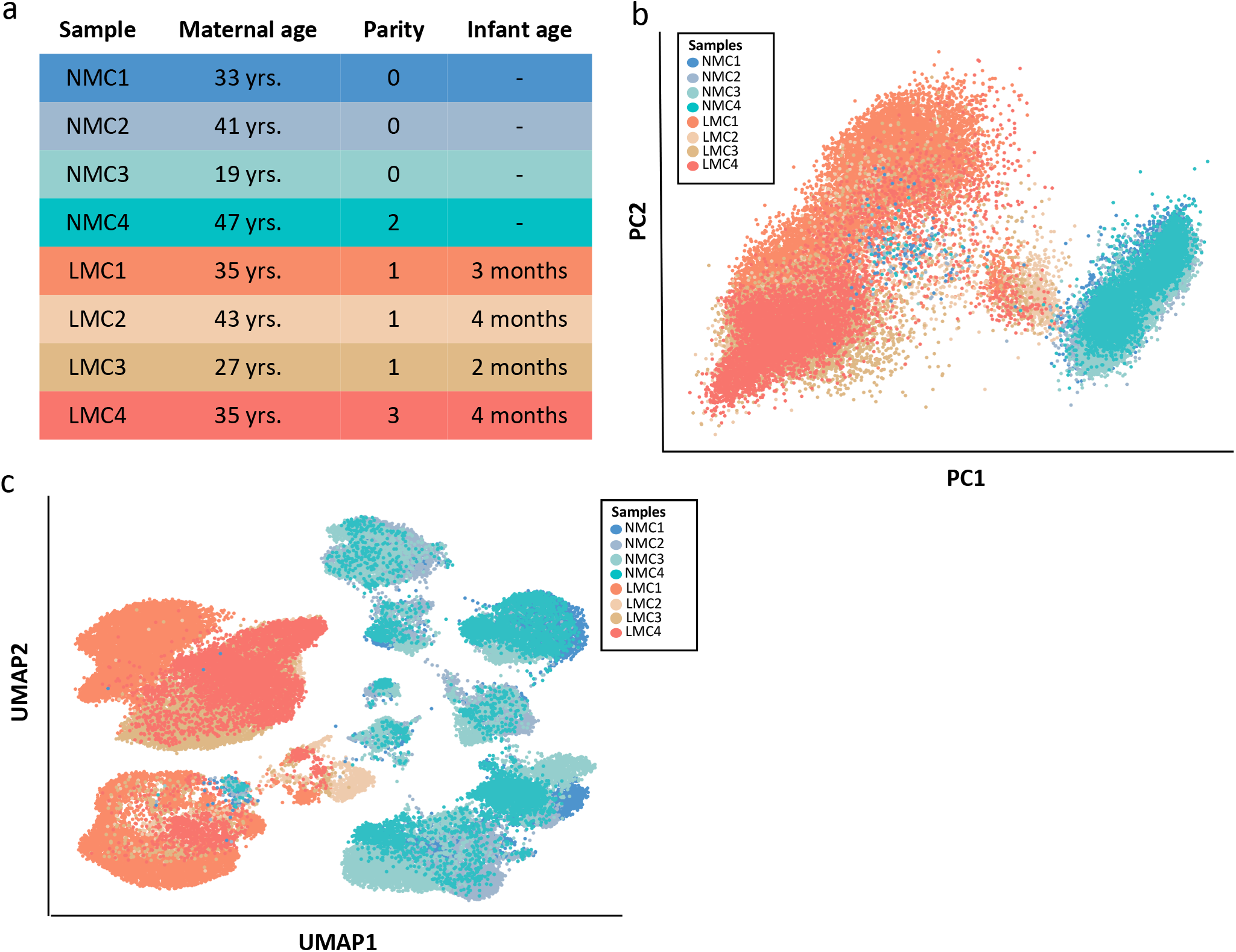
Understanding the differences between cells contributed by each non-lactating tissue (NMC) and lactation-derived milk cell (LMC) sample. **a)** Table describing the demographics of each participant **b)** Principal component (PC) analysis of all filtered and normalized cells revealed that the greatest variation along PC1 was due to samples coming from either NMCs or LMCs. **c)** Uniform manifold approximation and projection (UMAP) dimensional reduction of the mammary cells reveals distinct clusters arising from NMCs and LMCs where cells are colored by donor.

**Figure S3:**
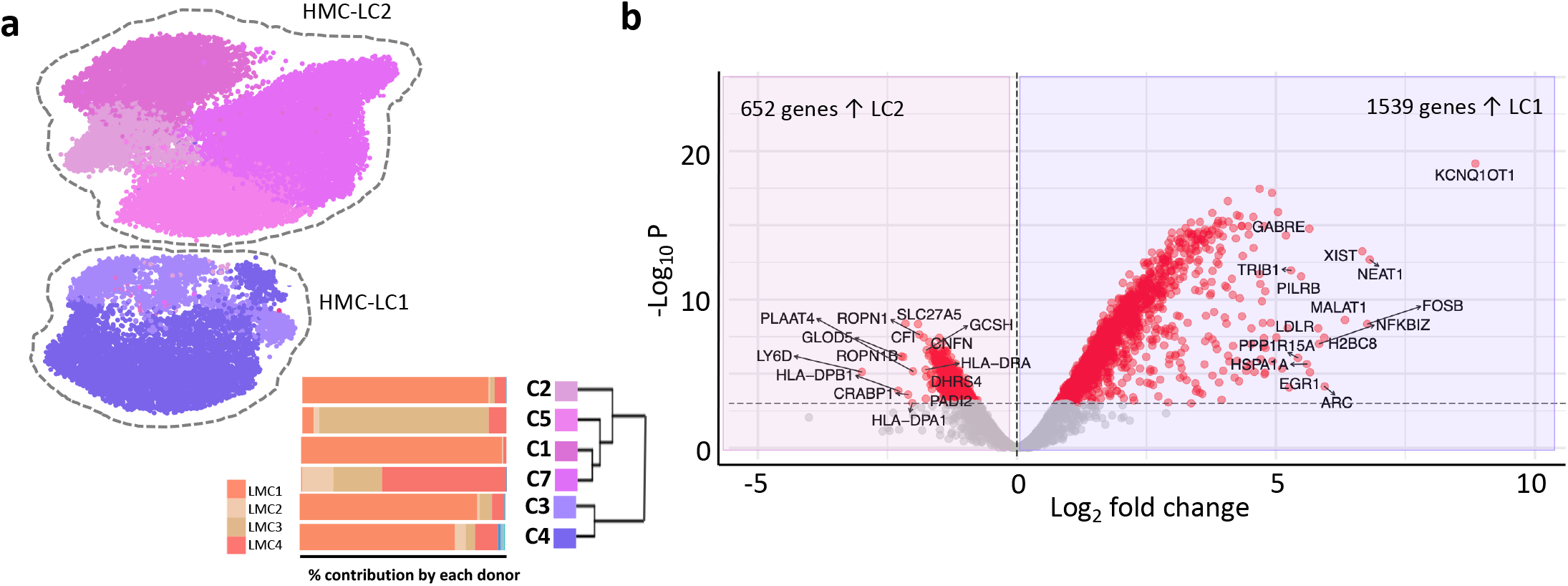
Exploring human milk lactation derived mammary cell (LMC) heterogeneity by comparing luminal clusters LC1 and LC2. **a)** Cellular contribution from each of the LMC sample donors to the luminal cell subclusters reveals unequal distribution **b)** Differential gene expression analysis revealed 652 genes highly expressed in LC2 compared to 1,539 genes more highly expressed in LC1 as displayed by a volcano plot (for a full list see Supplementary Table 1).

**Supplementary Figure 4:**
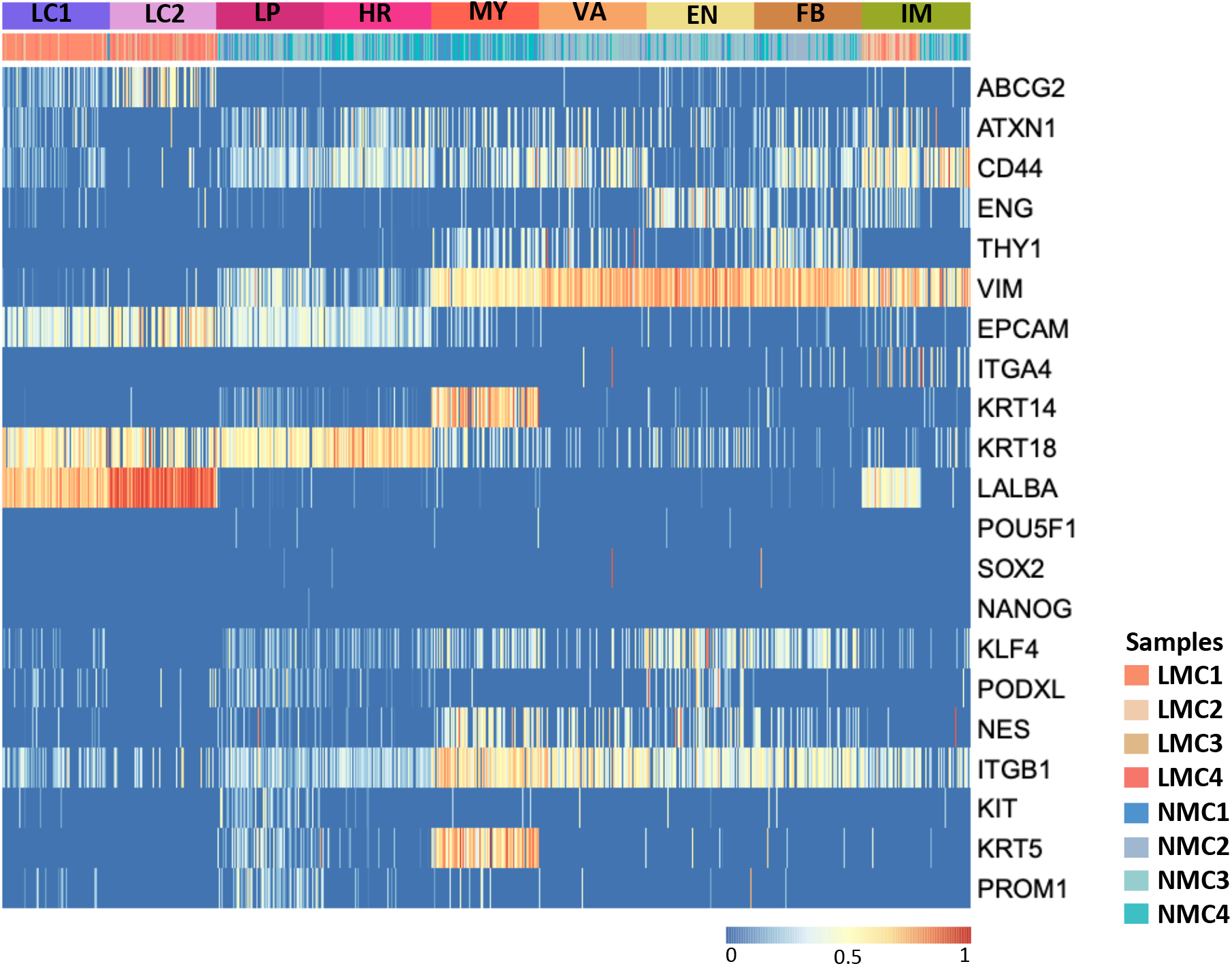
Heatmap displaying the expression of key genes previously described in human milk cells across both LMC and NMC clusters.

**Supplementary Figure 5:**
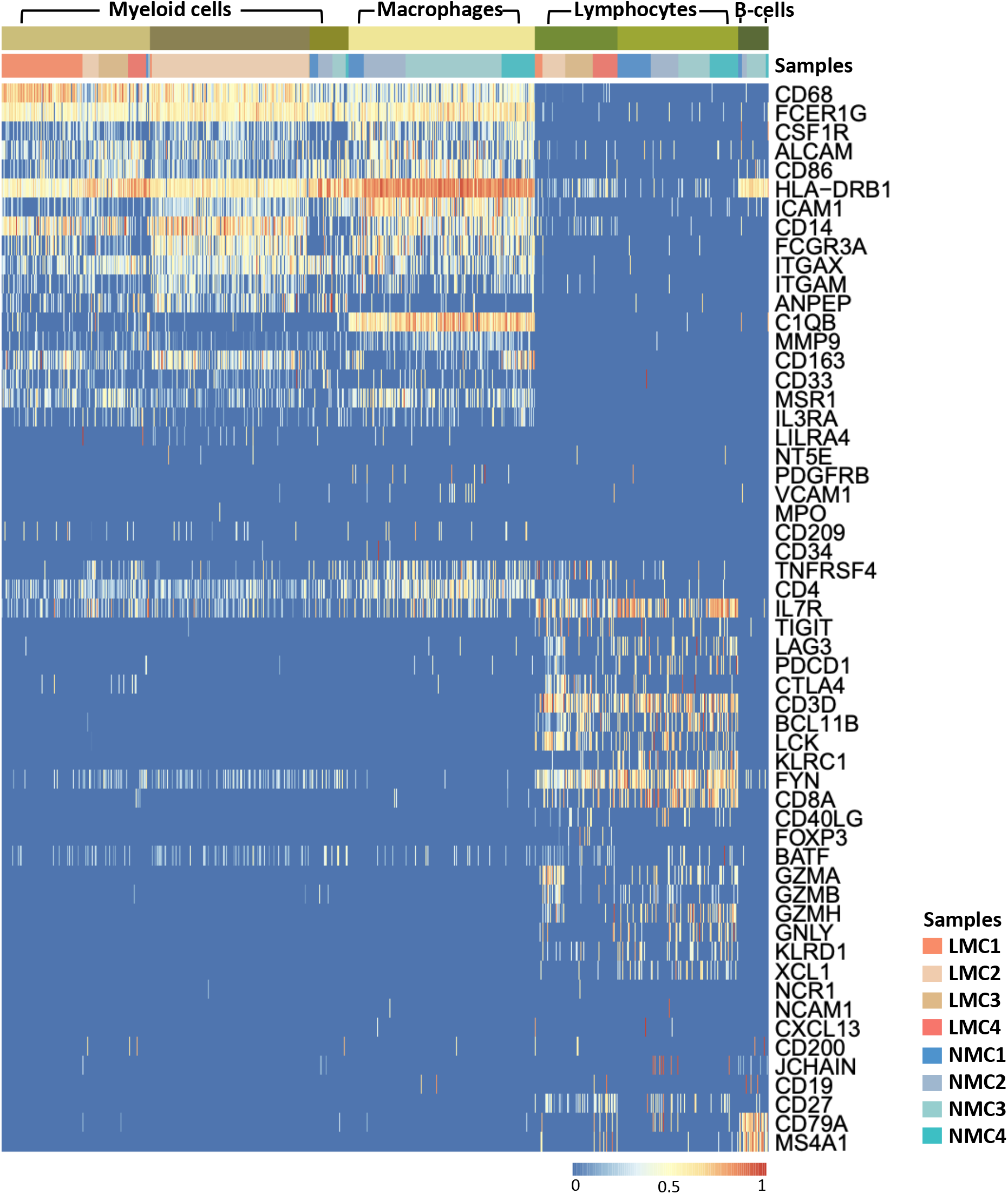
Heatmap displaying the expression of genes characteristic of different immune cell subpopulations across both LMC and NMCs.

**Supplementary Table 1:** DEGs between LC1 and LC2

**Supplementary Table 2:** GO terms for DEGs between LMC luminal cells, upregulated in LC2

**Supplementary Table 3:** GO terms for DEGs between LMC luminal cells, upregulated in LC1

**Supplementary Table 4:** Published mammary cell subpopulation scores

**Supplementary Table 5:** Differentially expressed genes between human milk luminal cells (LCs) and non-lactating Luminal Progenitors (LPs).

**Supplementary Table 6:** GO terms for DEGs between luminal cells which were upregulated in luminal LMCs

**Supplementary Table 7:** GO terms for DEGs between luminal cells which were upregulated in NMC LPs

